# A Predictive Model for Compound-Protein Interactions Based on Concatenated Vectorization

**DOI:** 10.1101/2024.10.02.616275

**Authors:** Gareth Williams, Kaz Azim

## Abstract

**Background:** Large data sets of compound activity lend themselves to building predictive models based on compound and target structure. The simplest representation of structure is via vectorisation. Compound fingerprint vectorisation has been successfully employed in predicting compound activity classes.

**Results:** A vector representation of a protein-compound pair based on a concatenation of the compound fingerprint and the protein triplet vector has been used to train random forest and neural network models on multiple datasets of protein-compound interaction together with compound associated transcription and activity profiles. Results for compound-target predictability are comparable with more complex published methodologies.

**Conclusion:** A simple intuitive representation of a protein-compound pair can be employed in a variety of machine learning models to gain a predictive handle on the activity of compounds for which there is no activity data. It is hoped that this transparent approach will prove sufficiently portable and simple to implement that drug discovery will be opened up to the wider research community.

## Introduction

There is a wealth of compound and protein interaction data deposited on platforms such as ChEMBL[1], with activity data on 2.4 million compounds and over 15 thousand targets, BindingDB[2], with 1.3 million compounds and 9 thousand targets, DrugBank[3], with 29 thousand drug-target pairings. This has spawned multiple efforts to generate predictive models whereby novel compound activities against targets can be predicted[4-8]. The models are largely based on representing the compound and protein as vectors. The minimal unambiguous compound description is provided by a simplified molecular-input line-entry system (SMILES)[9]. A compound SMILES allows for the 2D chemical structure to be defined and as such SMILES can be used to populate a bit string, or fingerprint, where each element represents the presence of a given element or chemical sub-structure. One popular compound fingerprinting is provided by PubChem[10] (https://ftp.ncbi.nlm.nih.gov/pubchem/specifications/pubchem_fingerprints.pdf), consisting of 881 structure elements. There are other fingerprinting schemes capturing differing levels of detail. Such as, the 166 element Molecular ACCess System (MACCS) keys[11] and the 1024 bit Extended Connectivity Fingerprint (ECFP)[12]. Fingerprinting facilitates fast compound similarity searching through a simple Tanimoto overlap scoring and have been used for screening large compound libraries[11, 13, 14]. This vectorisation of compound structure has been used to build a Bayesian model to effectively segregate compounds according to blood brain barrier penetration[15]. More recently, fingerprinting has been successfully employed to train a neural network on compound antibiotic activity leading and the repurposing of the experimental antidiabetic Jun kinase inhibitor halicin as a broad spectrum antibacterial[16]. Interestingly, halicin also emerges a s potential candidate antibiotic based on a simpler decision tree analysis of the training data[17].

In the present work we present a predictive framework for compound-protein interaction prediction based on representing the protein-compound pair as a simple concatenation of their vector representations. The compound vector is defined by the corresponding fingerprint and the protein is encoded by a 8,000 bit vector of amino acid triplets. We show that a basic random forest model is sufficient to recapitulate the level of predictability reported across a range of datasets. We apply this methodology to a dataset of IC50 data for compound-protein interactions gathered from ChEMBL, show that the model attains good predictability, with an area under the precision recall curve (AUPR) of 0.907+/-0.002.

In addition to compound activity data relative to specific targets, there is a wealth of publicly available high content quantitative activity data. The most extensive of these data comes from gene expression profiling through microarray and RNAseq platforms, deposited on the NCBI Gene Expression Omnibus (GEO)[18] and the EBI ArrayExpress[19] portals. Gene expression has facilitated links between disease states and drug activities that have motivated drug repurposing initiatives[20-24]. In this context, it is of interest to see whether transcriptional activity can be modelled on compound structure content. Extensive data on compound transcriptional activities have been compiled as part of the Broad Connectivity Map (CMAP) and Library of Integrated Network-Based Cellular Signatures (LINCS) initiatives[25, 26]. The latter data is based on gene level analysis of 978 landmark genes from which full profiles are predicted. We sourced the landmark gene data from GEO and generated categorical profiles based on expression thresholds pooled from multiple cell lines and treatment times. We found that a simple concatenation of expression profile and compound structure fingerprint provides sufficient content to tarin random forest and neural network models to provide moderate predictability.

Finally, we sourced data on compound cell line growth inhibitory activity generated by the National Cancer Institute (NCI)[27]. The data consists of inhibition data on 60 cancer cell lines and we found that random forest modelling could effectively predict compound activities based on combining compound fingerprints with numerical activity profiles.

In conclusion, our simple concatenated vectorisation of compound structure and target provides an effective representation to power predictive model building through random forest and neural network algorithms thus facilitating drug repurposing and discovery initiatives. The analysis also provides a simple portable pipeline for machine learning based on widely used packages and thus open to the wider research community.

## Methods

### Chemical fingerprints

Compound SMILES were mapped to chemical fingerprints using the R packages rcdk[28], to parse SMILES strings, and the fingerprint package (cran.r-project.org/web/packages/fingerprint/index.html) to build the fingerprint strings based on the SMILES. Compounds were mapped to maccs (166 dimensions), pubchem (881 dimensions) and extended (1024 dimensions) fingerprints.

### Protein vectorisation

Each amino acid triplet in the protein is represented by an integer *r*_*i*_ + 20*r*_*i*+1_ + 20^2^*r*_*i*+2_, where *r*_*i*_ = 0, …, 19 represent the amino acid type at position *i* along the protein chain. The protein vector is a 20^3^ = 8000 dimensional vector *v*_*n*_ = 0,1 according to the absence or presence of the n^th^ triplet.

### Databases for replication

Existing sets of training and test collections of protein-compound pairs were downloaded from the ConPlex[4] github page (github.com/samsledje/ConPLex_dev/tree/main/dataset). These data were sourced from publicly available data on protein-ligand binding affinities (BindingDB)[2], drug-target pairs from the Stanford BioSNAP collection[29] and kinase inhibitors (DAVIS)[30]. The decoy, non-interacting pairings, and training/testing data splits we used were the same as for the ConPlex analysis. Models were trained on the training datasets and results reported for the test data.

### ChEMBL derived protein-compound interaction data

The ChEMBL compound activity against protein target data was extracted from the ChEMBL activities database. ChEMBL hosts 2,716,345 compound interactions with IC50 data. Of these, 1,251,604 are for single mammalian protein targets. The protein interaction data was downloaded and filtered, with interactions defined based on IC50 falling below 50nM. This left 2,449 protein targets and 195,406 compounds distributed over 243,042 interactions. Many of the compounds are structurally similar and to avoid bias in predictive model building and ensure that the training and testing datasets are populated with structurally different compounds we restricted our analysis to a reduced diverse set of compounds. Multiple diverse sets of compounds were generated based on starting with a random compound and filtering the dataset to exclude those compounds with a structural Tanimoto similarity of 0.8 or above, based on a maccs fingerprint representation of the compound SMILES code. A second compound is then selected from the remaining dataset and the iteration proceeds to the last compound in the dataset, we are left with diverse sets of 23,860+/-63 compounds. Datasets for predictive model building were generated based on randomly sampling up to 50 compounds per protein target and for each of the proteins in these pairs randomly picking a non-interacting compound as a decoy. This resulted in 41,187+/-210 match/mismatch protein-compound pairs.

### LINCS data set

LINCS data for the 978 landmark gene set were downloaded from NCBI gene expression omnibus (GEO) with series accessions GSE92742 and GSE70138. The data consists of expression data corresponding to 83 cell types treated with 21,280 compounds for a variety of concentrations and treatment times. Of these compounds 19,692 were mapped to SMILES codes. Log normalised expression data were linearly regressed against compound dose for each plate. Z scores for each compound were then pooled across cell types and treatment times leaving one profile per compound. Profiles were then discretised according to *Z* ≥ 2 → 1, 2 > *Z* > −2 → 0, *Z* ≤ −2 → −1. The data together with compound SMILES is given in Supplementary Table S1. For the predictive model building each compound was paired with its transcriptional profile and with a random decoy profile that was less than 0.75 similar by Tanimoto score. Training and testing datasets had no compounds in common.

### NCI growth inhibition data

The National Cancer Institute (NCI) hosts data on the growth inhibitory activity for compounds across 60 cancer cell lines. We found the SMILES codes for 47,153 compounds. Predictive modelling was based on the data with the addition of one random profile per compound as a decoy ensuring that the true and decoy profiles have an RMSD of above 1. The data together with compound SMILES is given in Supplementary Table S2.

### Prediction model generation

The concatenated compound-protein vectors and associated binary match labels, where matching means interaction, were generated for the various datasets. Random forest and neural network models were trained on the training subset of the data and tested on the test subsets. The models were built within the R environment and python scripting. We built random forest models using the randomForest package (cran.r-project.org/web/packages/randomForest/)[31] and the faster implementation ranger[32]. We found that results were broadly in agreement and based on the speed of implementation, we generated multiple run statistics with ranger and these are the results presented here. Neural network model building was performed with tensorflow[33]. Tenosorflow modelling can also be implemented in R (cran.r-project.org/web/packages/tensorflow/). Model predictability was assessed with receiver operating characteristic (ROC) area under the curve (AUC) and area under the precision recall curve (AUPR). Statistics were based on results from different random initiations of training runs, when analysis was based on deposited training/test data splits, and different training/test data splits with our own protein interaction datasets.

## Results

### Comparative analysis of the BindingDB, BioSNAP and DAVIS datasets

We sought to validate the methodology against the datasets analysed as part of the recently published ConPlex predictive scheme. Here, three sets of data corresponding to protein-compound pairings were analysed, as described in Methods. Using the same training and testing subsets of the data that were deposited as part of the ConPlex study we trained a random forest on the training set and tested this on the validation set. The results for various compound fingerprint mappings are given in Table 1. As can be seen the predictability is enhanced with fingerprints containing a larger number of features. Our results compare favourably with those reported as part of the ConPlex study (BindingDB 0.628 ± 0.012, BioSNAP 0.897 ± 0.001 and Davis 0.458 ± 0.016). In addition to modelling the data using random forests we also trained a fast forward sequential neural network configured with two hidden layers of 20 nodes each and running for 200 epochs. We found that the neural network underperforms relative to the random forest, at least with settings that we tried, see Table 2.

**Table 1.** The predictive performance of a random forest model for the BindingDB, BioSNAP and DAVIS datasets. The predictive performance is measured through ROC AUC and AUPR from five independent runs corresponding to different random tree compositions of the forest. Predictability is enhanced with compound fingerprints encoding more structural information.

| BindingDB | ROC | AUPR | accuracy | sensitivity | specificity |
| --- | --- | --- | --- | --- | --- |
| maccs | 0.896+/-0.001 | 0.629+/-0.002 | 0.811+/-0.001 | 0.420+/-0.001 | 0.967+/-0.001 |
| pubchem | 0.913+/-0.000 | 0.674+/-0.001 | 0.831+/-0.001 | 0.452+/-0.001 | 0.970+/-0.001 |
| extended | 0.920+/-0.001 | 0.692+/-0.001 | 0.840+/-0.002 | 0.468+/-0.003 | 0.970+/-0.001 |
| BIO SNAP | ROC | AUPR | accuracy | sensitivity | specificity |
| maccs | 0.869+/-0.001 | 0.892+/-0.000 | 0.805+/-0.002 | 0.834+/-0.001 | 0.781+/-0.002 |
| pubchem | 0.885+/-0.001 | 0.904+/-0.001 | 0.815+/-0.002 | 0.838+/-0.002 | 0.794+/-0.002 |
| extended | 0.885+/-0.001 | 0.906+/-0.001 | 0.820+/-0.001 | 0.852+/-0.001 | 0.793+/-0.001 |
| DAVIS | ROC | AUPR | accuracy | sensitivity | specificity |
| maccs | 0.853+/-0.005 | 0.359+/-0.010 | 0.798+/-0.002 | 0.164+/-0.002 | 0.983+/-0.000 |
| pubchem | 0.900+/-0.001 | 0.467+/-0.010 | 0.848+/-0.002 | 0.219+/-0.002 | 0.987+/-0.001 |
| extended | 0.911+/-0.002 | 0.479+/-0.009 | 0.851+/-0.003 | 0.225+/-0.003 | 0.988+/-0.000 |

**Table 2.** The predictive performance of a standard neural network with two 20 node hidden layers model for the BindingDB, BioSNAP and DAVIS datasets. The predictive performance is measured through ROC AUC and AUPR from five independent runs corresponding to different random tree compositions of the forest. Predictability is again enhanced with compound fingerprints encoding more structural information.

| **BindingDB** | **ROC** | **AUPR** | **accuracy** | **sensitivity** | **specificity** |
| --- | --- | --- | --- | --- | --- |
| **maccs** | 0.877 +/- 0.003 | 0.492 +/- 0.007 | 0.781 +/- 0.005 | 0.856 +/- 0.009 | 0.768 +/- 0.007 |
| **pubchem** | 0.886 +/- 0.002 | 0.521 +/- 0.008 | 0.796 +/- 0.004 | 0.854 +/- 0.006 | 0.786 +/- 0.006 |
| **extended** | 0.898 +/- 0.002 | 0.551 +/- 0.002 | 0.825 +/- 0.004 | 0.838 +/- 0.016 | 0.823 +/- 0.008 |

| **BIO SNAP** | **ROC** | **AUPR** | **accuracy** | **sensitivity** | **specificity** |
| --- | --- | --- | --- | --- | --- |
| **maccs** | 0.856 +/- 0.005 | 0.849 +/- 0.005 | 0.784 +/- 0.007 | 0.804 +/- 0.016 | 0.763 +/- 0.020 |
| **pubchem** | 0.863 +/- 0.004 | 0.855 +/- 0.006 | 0.792 +/- 0.005 | 0.800 +/- 0.004 | 0.785 +/- 0.014 |
| **extended** | 0.883 +/- 0.004 | 0.873 +/- 0.005 | 0.813 +/- 0.003 | 0.817 +/- 0.009 | 0.809 +/- 0.010 |

| **DAVIS** | **ROC** | **AUPR** | **accuracy** | **sensitivity** | **specificity** |
| --- | --- | --- | --- | --- | --- |
| **maccs** | 0.826 +/- 0.008 | 0.170 +/- 0.005 | 0.749 +/- 0.012 | 0.798 +/- 0.018 | 0.746 +/- 0.014 |
| **pubchem** | 0.870 +/- 0.007 | 0.244 +/- 0.012 | 0.813 +/- 0.007 | 0.782 +/- 0.017 | 0.814 +/- 0.008 |
| **extended** | 0.893 +/- 0.003 | 0.296 +/- 0.021 | 0.810 +/- 0.007 | 0.843 +/- 0.020 | 0.808 +/- 0.008 |

### ChEMBL derived protein-compound interaction data

ChEMBL compound IC50 interaction data with proteins as targets was processed into training and testing datasets as described in the Methods section. The results are broadly in agreement with the BioSNAP and BindingDB analyses, see Figure 1.

**Figure 1.**
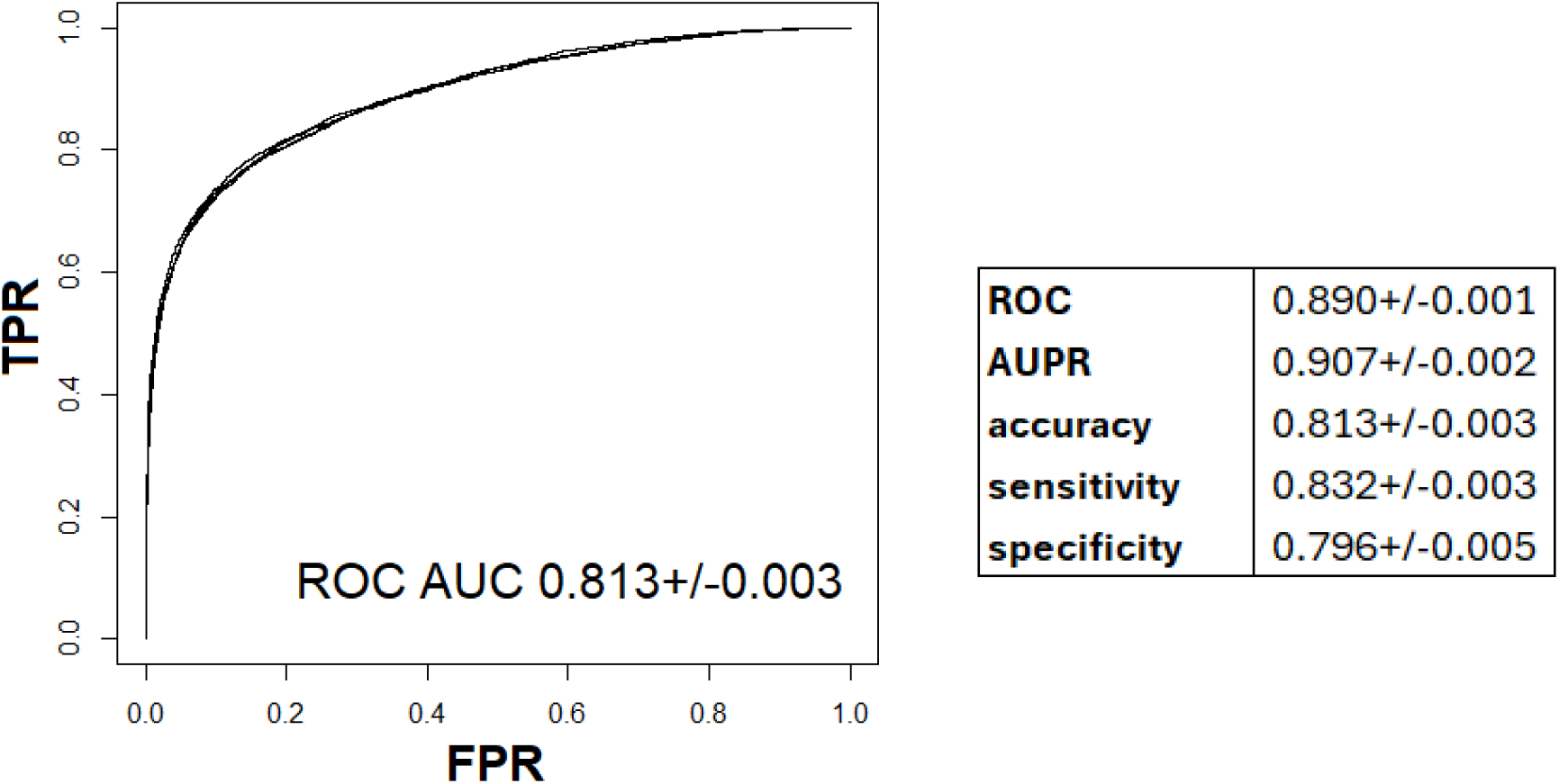
Random forest model results for predicting the protein target inhibited by a compound. The ROC curves for four independent runs based on different diverse compound sets are shown to the left and the statistics are given in the table at the right. The data was based on compound inhibition data extracted from ChEMBL. The predictability is broadly in line with that for the BioSNAP and BindingDB data analysed in the previous section.

The random forest modelling has the potential to correctly assign protein target to compounds with no structural similarity to training set compounds. In Figure 2 four compounds with correct protein target assignments are shown. These compounds had no structurally similar compounds in the training set targeting the same protein. The training set compound shown is the most structurally similar to the testing set compound against the same target protein.

**Figure 2.**
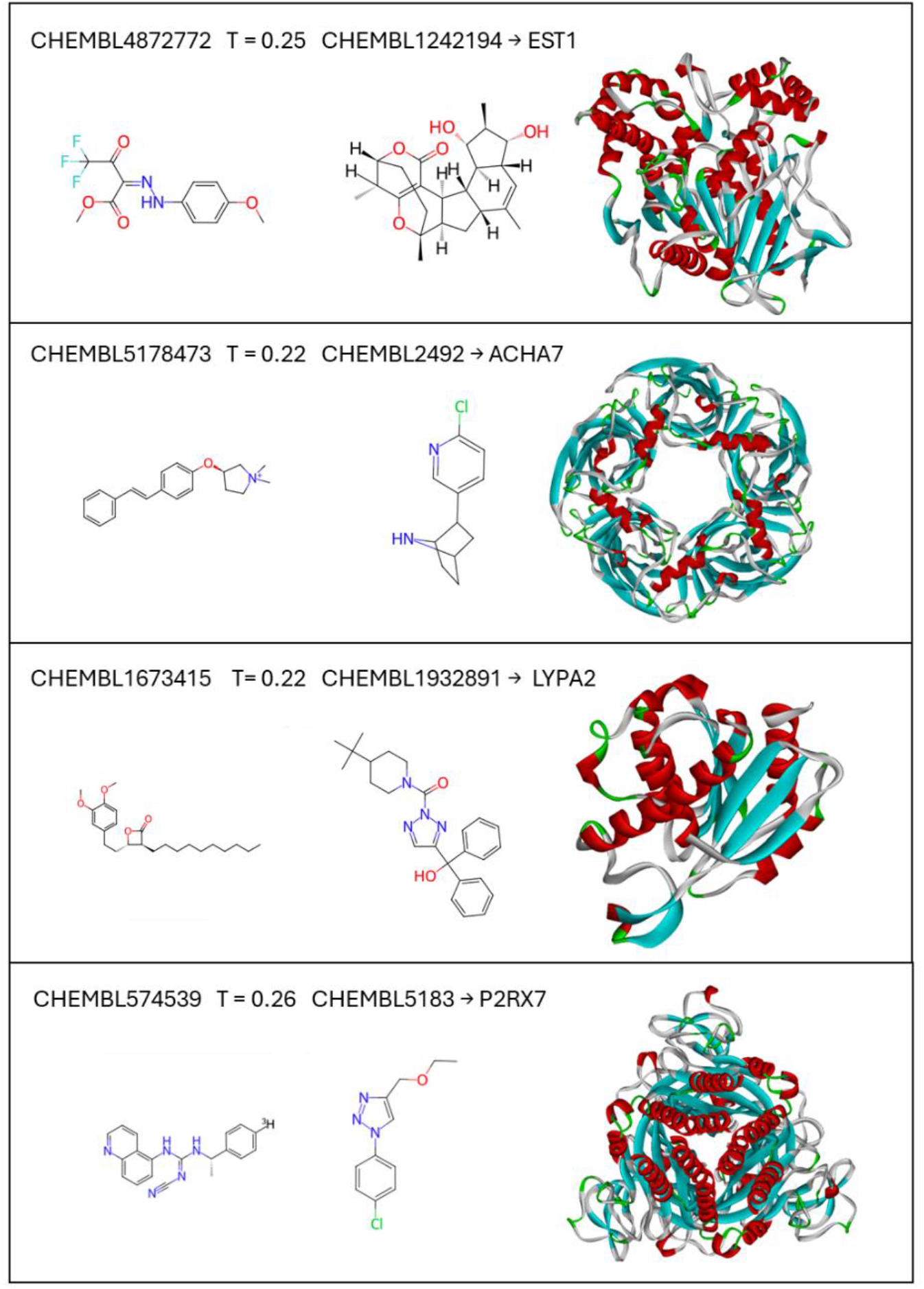
The random forest model correctly assigns activities to novel structures. Four pairs of structurally dissimilar compounds with maccs fingerprint Tanimoto scores shown together with protein target. The compound on the left was correctly predicted to target the protein show. The compound on the right was the most structurally similar compound in the training set targeting this same protein.

### Predicting compound transcription profiles

Next, the potential of this simple vectorisation approach to model the associations of compound structure with activity profiles was examined. Gene expression provides for a high content quantitative phenotypic readout of compound activity and there is plentiful data in the public domain, see Introduction. As described in the Methods section, we compiled expression change thresholded categorical compound profiles pooled from multiple cell line and treatment time data from the LINCS landmark expression data. The data was supplemented with decoy mismatched compound-profile pairs and split into training and testing subsets. Training a random forest model, using the ranger package in the R environment, on the training data set we find moderate predictability within the testing subset, see Table 3. Interestingly, a neural network model appears to consistently outperform the random forest model, see Table 3 under NN. It would be of interest to see what underlies this enhanced performance of the neural network model, given that random forest modelling outperforms this approach in all the other datasets we have analysed for this study.

**Table 3.** Random forest and neural network models predict the association of compound to transcriptional profile based on the compound fingerprint. The results from multiple decoy sets are shown. The random forest results (RF) show decreased predictability relative to the neural network (NN) model.

|  | RF | NN |
| --- | --- | --- |
| **ROC** | 0.779+/-0.003 | 0.816+/-0.005 |
| **AUPR** | 0.738+/-0.001 | 0.771+/-0.008 |
| **accuracy** | 0.716+/-0.005 | 0.744+/-0.004 |
| **sensitivity** | 0.710+/-0.002 | 0.740+/-0.010 |
| **specificity** | 0.722+/-0.009 | 0.747+/-0.011 |

In effectiveness of the predictive model can be seen through the ability to predict the transcription profiles of compounds for which there are no similarly structured compounds in the data on which the model was trained, see Figure 3.

**Figure 3.**
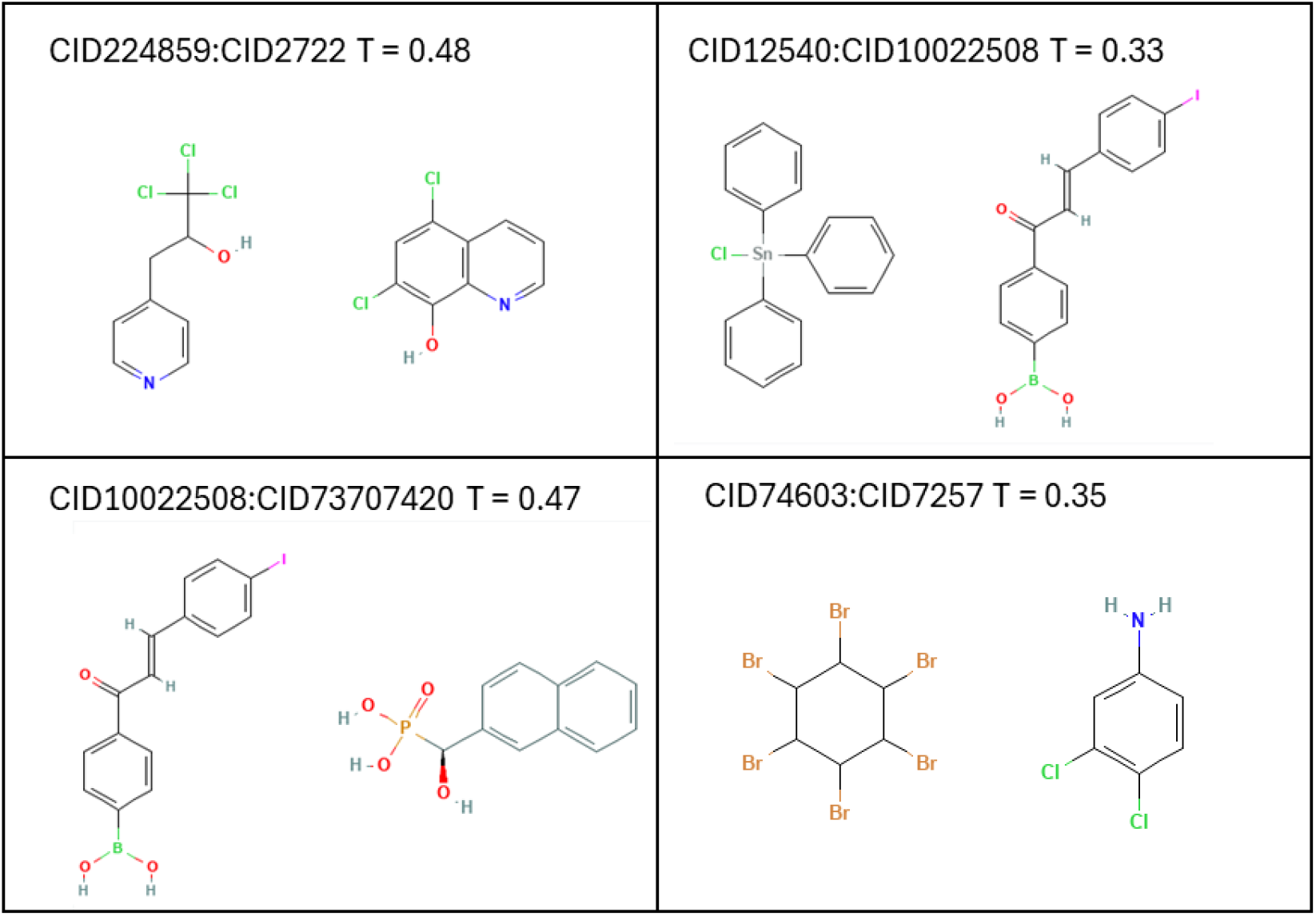
Transcriptional profiles can be assigned to compounds with novel structures. Example compounds for which the correct transcriptional profile was predicted shown together with the most similar compound in the whole of the training set (measured by maccs Tanimoto similarity). The compound on the right in the pair was part of the training set.

### Predicting compound cell-line growth inhibition

In the case of the NCI-60 data the activity profiles consist of numerical readouts across 60 cell lines as opposed to categorical transcription profiles used above. In this context, it is interesting to see to what extent the random forest approach can segregate matched compound profile pairs from decoy pairs. The level of predictability is slightly higher than for transcription profiling, see Table 4. This is somewhat surprising as the LINCS profiles are of higher dimensionality but may be due to the larger number of compounds in the NCI-60 dataset. As with the LINCS analysis it is clear that the modelling can predict the activities of compounds for which there are no structurally similar occurrences in the training data, see Figure 4.

**Table 4.** Random forest and neural network models predict the association of compound to NCI-60 profile based on the compound fingerprint. The results from multiple decoy sets are shown. The random forest results (RF) show significantly better predictability relative to the neural network (NN) model.

|  | RF | NN |
| --- | --- | --- |
| **ROC** | 0.787+/-0.003 | 0.649+/-0.015 |
| **AUPR** | 0.778+/-0.004 | 0.611+/-0.015 |
| **accuracy** | 0.707+/-0.004 | 0.612+/-0.013 |
| **sensitivity** | 0.691+/-0.004 | 0.691+/-0.053 |
| **specificity** | 0.727+/-0.003 | 0.532+/-0.043 |

**Figure 4.**
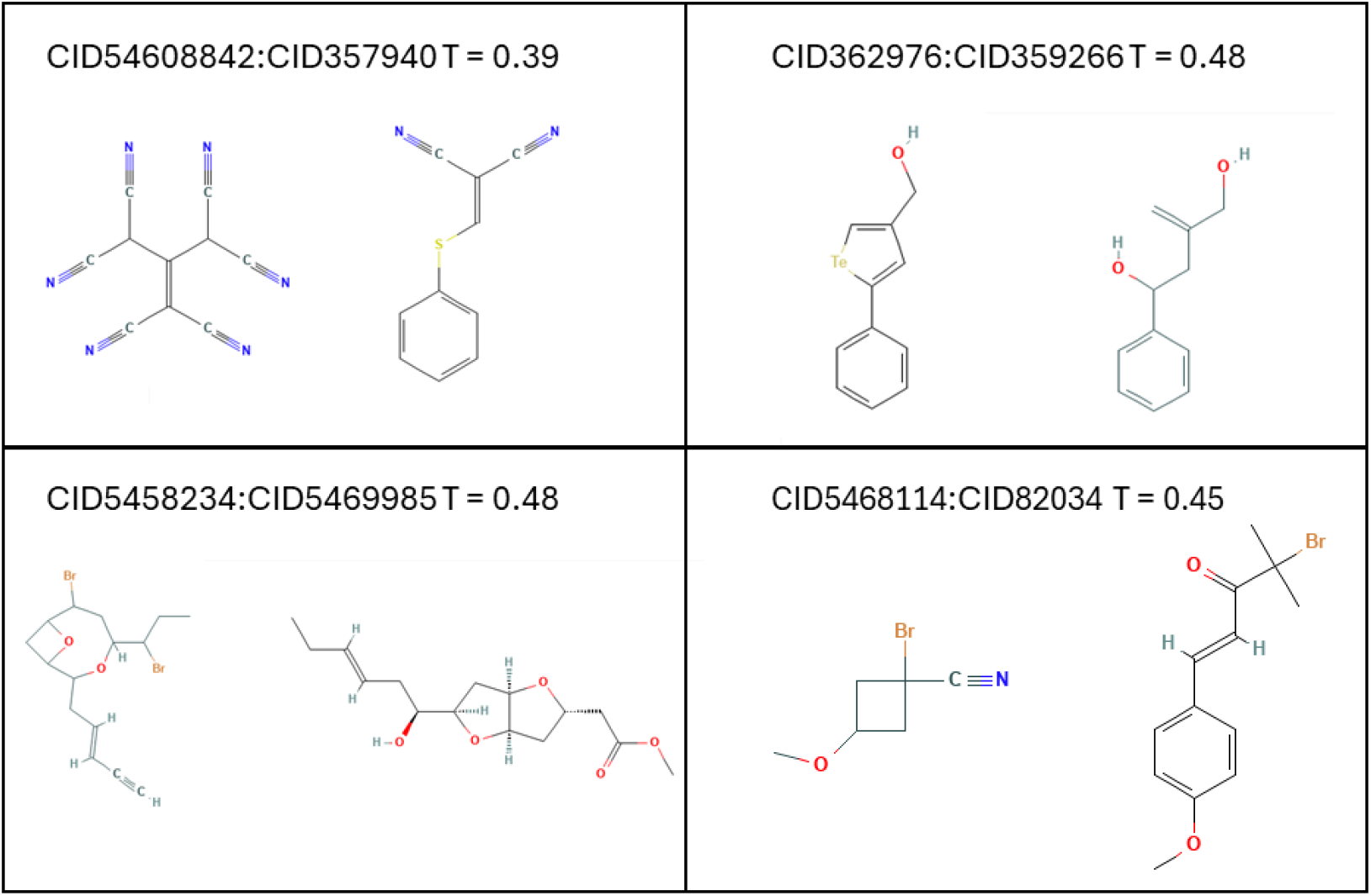
Correct inhibitory profiles can be assigned to compounds with novel structures. Example compounds for which the correct NCI-60 profile was predicted shown together with the most similar compound in the whole of the training set (measured by maccs Tanimoto similarity). The compound on the right in the pair was part of the training set.

## Conclusions

Concatenating vectors representing compound structure and activity, in the form of interaction target or high content activity data, facilitates training random forest and neural network models to gain a predictive handle on the activity of novel compounds. We have found that this simple methodology performs as well as published algorithms in predicting protein targets in three distinct databases that cover various aspects of compound activity. We applied the methodology to large compound interaction datasets extracted from ChEMBL and found a consistently high level of predictability with a ROC AUC of 0.890+/-0.001 and an AUPR of 0.907+/-0.002. The model was able to assign the correct protein target to compounds sharing no structural similarity with the compound set on which the model was trained. In addition, we applied the methodology to compound activity profiles in the form of transcriptional data and inhibitory activity against a set of cell lines. We found that the simple vectorisation approach returns reasonable predictability. With a wider repertoire of ChEMBL target associations it is hoped that an application of the methodology could be to de-orphan drugs, with query drugs compared to drugs with established activities based on correlating the predicted interaction scores against the expanded protein target set. It would also be of interest to see to what extent additional content in protein vectorisation will impact on predictability. Protein vectorisation could for example include information about the presence/absence of a set of well defiled motifs. Another avenue that could be pursued would be to hash longer strings of amino acids and keeping the vectors within reasonable bounds by mapping amino acids into a reduced set of categories based on shared physicochemical properties or mutual substitutional rates.

In conclusion, we have presented a simple and portable methodology for predicting the activities of compounds based on combining the compound fingerprint and a vector encoding compound activity, either in the form of the protein target or multidimensional biological readout. The method compares favourably when tested against published methodologies and is simple to implement. It is hoped that this will allow for researchers to harness the wealth of publicly available compound activity data with a view to speeding up drug repurposing.

All scripts used as part of the study are available through the github at https://github.com/GarethWgh/CIDpred. All enquiries to GW.

## Notes

### Competing Interest Statement

The authors have declared no competing interest.

